# Optimized In Vitro Expansion of Vδ1+ γδT Cells from Nonhuman Primate Peripheral Blood

**DOI:** 10.64898/2025.12.03.692135

**Authors:** Isabelle Berthelot, Alexandra McNally, Victoria Hart, Yoshinori Fukazawa, Nizar Batada, Namita Rout

## Abstract

Gamma delta (γδ) T cells are a subset of T cells that express MHC-independent γδ T cell receptors (TCRs) and can perform the same T-helper functions as CD4+ cells as well as cytotoxic functions like CD8+ T cells. The MHC-independent nature of γδ TCRs allows them to recognize a larger diversity of antigens, including self and non-self-antigens, making them ideal candidates for immunotherapeutic interventions. Both translational science efforts to better understand the physiology of γδ T cells and clinical research concerning the applications of γδ T cell immunotherapy require successful γδ T cell expansion techniques. Building on prior methods and recent innovations, in this study we optimized in vitro methods for rapid and efficient expansion of Vδ1^+^ T cells from stimulated PBMCs. CD3-stimulated PBMCs from rhesus macaques in the presence of phospho-vitamin C (pVC) and IL-15 achieved up to a 6561-fold expansion (average 2559-fold) increase of Vδ1^+^ T cells after 9-day culture. Comparable expansion was obtained with phytohemagglutinin (PHA)-stimulated PBMCs, achieving up to 7574-fold expansion (average 2040-fold) increase when IL-7 and IL-18 were added alongside IL-15 and pVC. Notably, inclusion of pVC significantly enhanced the expansion of Vδ1^+^ T cells in both stimulation conditions. These results provide optimized conditions for scalable in vitro expansion of peripheral blood Vδ1^+^ T cells from nonhuman primate models, supporting downstream applications in immunophenotyping, functional assays, and preclinical modeling of γδ T cell–based immunotherapies.

## Introduction

T cells play a vital role in the adaptive immune system. Since their discovery, most studies have focused on conventional αβ T cells, whose α and β T cell receptor (TCR) chains recognize peptide antigens presented by major histocompatibility complex (MHC) molecules. In the mid 1980s, a distinct lineage of T cells expressing γδ TCR was discovered^1, 2^. While γδ T cells can perform T-helper functions as CD4^+^ T cells as well as cytotoxic functions of CD8^+^ T cells, they are unique in their ability to recognize a larger range of antigens that are not presented on MHC molecules^3, 4^. Each γδ TCR is comprised of a pair of γ and δ chain, producing a wide range of TCR subtypes with each combination yielding different capabilities and residing in different areas of the body ^2^. The most widely studied γδ T cell subtypes are Vδ1^+^ and Vδ2^+^ T cells, which reside mainly in tissues and blood, respectively ^5^.

The MHC-independent nature of γδ TCRs allows them to recognize a larger diversity of self and non-self antigens, making them ideal candidates in immunotherapeutic research^2, 5, 6^. Adoptive transfer of allogeneic immune cells is made possible by γδ T cells, opening a new technology for immunotherapy that cannot be achieved by conventional αβ cells due to risk of graft-versus-host disease ^5^. Among γδ T cell subsets, Vδ2^+^ T cells are the predominant population in the peripheral blood of healthy individuals, providing a readily accessible source for in vitro expansion and functional studies ^7^. The Vδ2^+^ subsets have demonstrated robust tumor recognition and cytotoxic activity and are currently being investigated as a therapy for multiple malignancies, including head and neck cancers, hepatocellular carcinoma, and prostate cancer ^4, 5^. In contrast, Vδ1^+^ T cells, though less abundant in circulation and less extensively characterized, exhibit potent tumor-killing activity and preferentially localize to mucosal and epithelial tissues, where they interact with target cells through their tissue-homing properties^2, 7, 8^. In a study that compared Vδ1^+^ versus Vδ2^+^ T cell subsets’ ability to infiltrate and kill human tumor cells injected into mouse model, it was found that Vδ1^+^ T cells showed increased effectiveness in tumor suppression and cytotoxicity, underscoring the need for optimized methods to expand and study this clinically relevant subset^4, 9^.

Although the potential of γδ T cells is increasingly recognized, they remain an understudied subset of immune cells. Roadblocks in the advances of translational γδ T cell research include developmental and functional differences in γδ T cells between rodents and humans, limiting the translational and clinical understanding of γδ T cell immunology ^4^. Murine γδ T cells are primarily defined by usage of distinct γ chains, while human γδ T cells are mainly categorized based on δ chain composition, complicating the extrapolation of murine findings to human biology ^6^. Moreover, the strong evolutionary divergence in TCRγ and TCRδ genes between rodents and primates underscores the need for primate models to accurately investigate γδ T cell physiology relevant to human immune response ^4^. In addition to difficulties in observing primate γδ T cell physiology in rodent models, the small population of γδ T cells in the blood and tissues hinder accessibility. Although frequencies vary depending on animal species and state of health, γδ T cells make up <1-5% of peripheral blood mononuclear cells (PBMCs), representing a minor fraction of circulating lymphocytes, with the Vδ1^+^ subsets being even rarer^5, 7^. The limited accessibility of γδ T cells for experimental studies is further compounded by the scarcity of commercial antibodies targeting γδ T cell-specific markers ^5^.

γδ T cell-based immunotherapy relies on the efficient isolation and expansion of the γδ T cell subsets from PBMCs for downstream manipulation or therapeutic reintroduction. Robust and reproducible expansion protocols are essential not only for proof-of-concept clinical studies but also for advancing translational research into γδ T cell biology ^5^. Most current expansion strategies are tailored for the expansion of Vδ2^+^ T cell subsets, which responds to aminobisphosphonates and phosphoantigens that drive selective in vitro expansion ^7^. In contrast, there are currently no validated agonists for in vivo expansion of Vδ1^+^ T cells, restricting their therapeutic use to in vitro culture systems ^4^, approaches that still require further optimization.

The ‘DOT Protocol’ ^9^ enabled robust expansion of cytotoxic Vδ1^+^ T cells, and subsequent studies have demonstrated that anti-CD3/IL-15 stimulation or phytohemagglutinin (PHA), a plant-derived mitogen used to induce T cell activation, with feeder cells can further optimize expansion depending on culture conditions^7, 8^. Despite these advances, important limitations remain, including challenges in scalability, reproducibility, and adaptation to preclinical models. In particular, optimized protocols for nonhuman primate (NHP) Vδ1^+^ T cells are lacking, restricting proof-of-concept studies necessary to evaluate γδ T cell-based therapies in a translational context. In a 2019 study by Kouakanou and colleagues demonstrated a beneficial effect of Vitamin C addition on γδ T cell proliferation and viability ^10^. This study optimized concentrations of both forms of Vitamin C, indicating that L-ascorbic acid (Vitamin C) optimized cell growth at 70 μM (12.5 μg/mL), while the more stable and less toxic phospho-Vitamin C (pVC) optimized cell growth at 173 μM (50 μg/mL). PVC-stimulated γδ T cells showed increased Ki-67 expression, a marker of cycling cells, indicating the enhanced proliferative effect of pVC on γδ T cells is associated with cell cycle progression^10^. This application of pVC on γδ T cell expansion show promise but has only been optimized for Vδ2^+^ T cells, not Vδ1^+^ T cells.

Here, we present an optimized method for the expansion of NHP Vδ1^+^ T cells that achieves up to 6561-fold increase, with an average of 2559-fold increase from cryopreserved PBMCs after 9 days of in vitro culture following TCR stimulation. Comparable Vδ1^+^ T cell expansions were obtained with phytohemagglutinin (PHA)-stimulated PBMCs, achieving up to 7574-fold expansion (averaging 2040-fold increase) when IL-7 and IL-18 were added alongside IL-15, and pVC. Moreover, we explore the effects of pVC supplementation on the proliferation of Vδ1^+^ T cells under various expansion conditions.

## Methods

### PBMC Isolation

Blood samples from healthy, uninfected Indian-origin rhesus macaques (*Macaca mulatta)* housed at the Tulane National Biomedical Research center were collected in EDTA vacutainer tubes (Sarstedt Inc., Order# 01.1605.100) and processed immediately. PBMCs were separated by density gradient centrifugation (Lymphocyte Separation Medium; MP Biomedicals, Cat# 50494X) at 1500 rpm for 45 minutes. PBMCs were suspended in Bambanker freezing media (Bulldog Bio, Cat# BB05) and cryopreserved in liquid nitrogen system at –180°C. All expansions were conducted from cryopreserved PBMCs.

### Cytokines and Reagents

Cytokines and reagents used for expansion cocktails include recombinant human IL-7 protein, Gibco™ human IL-18BP Fc recombinant protein, human IL-2 IS premium grade, recombinant human IL-15 carrier-free, phospho-Vitamin C, purified PHA, anti-CD3 antibody, and anti-CD28 antibody (Manufacturer and clone information found in Supplementary Table 1).

### In vitro Vδ1^+^ T cell expansion

Macaque PBMCs were stimulated using plate-bound anti-CD3 antibody or PHA in the presence of different cytokine combinations to induce Vδ1^+^ T cell expansion (Fig. 1). In all conditions 1×10^6^ PBMCs were co-cultured with irradiated human PBMC feeder cells at 1:2 ratio in 48-well plates in a final volume of 1 mL. Feeder PBMCs were exposed to 300 rads of γ-irradiation for 22 minutes or, alternatively, obtained pre-irradiated (iQ Biosciences, Cat# NC1863492). For plate-bound anti-CD3 stimulation, the wells were treated with anti-CD3 antibody and incubated for 2 hours at 37°C and aspirated after incubation, followed by seeding with 1×10^6^ PMBCs per well along with 2×10^6^ irradiated feeder cells. Six different cytokine stimulations were tested (Table 1, concentrations found in Supplementary Table 2). The cells were cultured in RPMI medium (Gibco, Cat# 11875119) supplemented with 10% FBS (Atlas Biologicals, Cat# F-0500-DR), 100 U/mL penicillin, and 100 μg/mL streptomycin. Expanding cells were split into 2–3 wells every 2–3 days in and replenished with fresh medium containing cytokines. The cells were cultured until day 9 and harvested for cell counts and flow cytometry. Fold expansion was calculated by dividing the final number of total and Vδ1^+^ T cells by their initial number on day 0, combining data from viability and cell counts as well as flow cytometric analysis.

**Table 1.** Stimulation conditions and cytokines used for γδ T cell expansion from PBMCs.

| Cocktail | Stimulation | Cytokines | pVC |
| --- | --- | --- | --- |
| A | PHA | IL-7, IL-18, IL-15 | - |
| B | PHA | IL-7, IL-18, IL-15 | + |
| C | TCR (plate-bound α-CD3 + soluble α-CD28) | IL-2, IL-15 | - |
| D | TCR (plate-bound α-CD3 + soluble α-CD28) | IL-15 | - |
| E | TCR (plate-bound α-CD3 + soluble α-CD28) | IL-15 | + |
| F | TCR (plate-bound α-CD3 only) | IL-15 | + |

**FIGURE 1.**
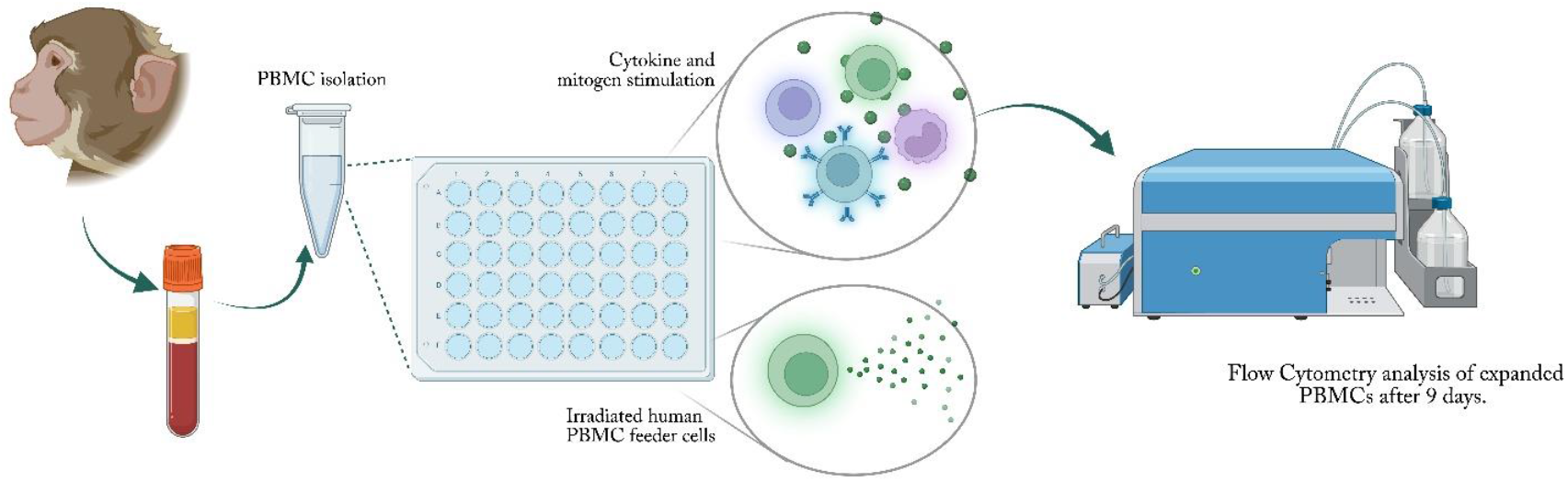
Study design: Macaque PBMCs were stimulated with plate-bound anti-CD3 or PHA and co-cultured with irradiated feeder PBMCs in the presence of various cytokine combinations. Cell cultures were harvested on day 9 for final cell counts, viability and immunophenotyping analyses.

### Flow Cytometry

γδ T cell subpopulation phenotype and purity were assessed using antibodies for CD45, CD3, CD4, CD8, TCR γδ, TCR Vδ1, and TCR Vδ2. Memory phenotype was assessed using CD95 and CD28 antibodies, and functional phenotype was assessed using antibodies for the transcription factors RORγt and T-bet (Manufacturer and clone information found in Supplementary Table 3). Expanded PBMCs were washed with PBS containing 2% FBS and counted prior to staining. Fixable Viability Stain (FVS700) was used for exclusion of dead cells from the analyses. Surface staining for extracellular markers was carried out for 30 minutes in 4°C. Intracellular staining for RORγt and T-bet was carried out using the BD Cytofix/Cytoperm fixation and permeabilization kit (BD, Cat# 554714), according to the manufacturer’s protocol. Flow cytometric analysis was conducted using BD Symphony A5 and BD Fortessa flow cytometers, and data were acquired with BD FACSDiva Software v9.3.1. Post-acquisition data processing was carried out using FlowJo software version 10. Gating strategy found in Supplementary Figure 1.

### Statistical Analysis

Statistical analyses were performed using GraphPad Prism version 10 (GraphPad Software). Data are presented as mean ± SEM. Statistical significance was determined using paired t-tests or Mann-Whitney test when applicable, with p < 0.05 (*), p < 0.01 (**), and p < 0.001 (***) considered significant.

## Results

### CD3/IL-15/pVC stimulation drives preferential and high-fold expansion of Vδ1^+^ T cells from PBMCs

Studies have shown that Vδ1+ T cells respond well to CD3 stimulation for in vitro expansion ^8, 11^, while others have shown efficient expansion following stimulation with mitogens such as ConA and PHA ^9, 12^. Thus, we tested various cytokine combinations with TCR (anti-CD3+anti-CD28) and PHA stimulations (Table 1) and assessed the expansion of Vδ1+ and Vδ2+ T subsets from PBMCs (Fig. 1 & 2A). Results from 9-day in vitro expansion under various conditions were evaluated based on the total number of Vδ1^+^ and Vδ2^+^ T cells obtained post-expansion (Fig. 2B) and their corresponding fold increases relative to baseline (Fig. 2C). The cell yields for each cytokine condition are summarized in Table 2 (Supplemental expansion results data found in Supplementary Table 4).

**Table 2.** Cell yields and fold-changes for Vδ1^+^ and Vδ2^+^ T cells from various stimulation conditions and cytokines used for γδ T cell expansion from 1 million PBMCs.

| Stimulation | Cocktail | Max Vδ1<sup>+</sup> yield | Avg. Vδ1<sup>+</sup> Fold Change | Min, max Vδ1<sup>+</sup> Fold Change | Max Vδ2<sup>+</sup> yield | Avg. Vδ2<sup>+</sup> Fold Change | Min, max Vδ2<sup>+</sup> Fold Change |
| --- | --- | --- | --- | --- | --- | --- | --- |
| PHA | A | 0.765e6 | 505 | 0.2, 2250 | 0.088e6 | 12 | 0.04, 46 |
| PHA | B | 2.88e6 | 2040 | 196, 7574 | 0.985e6 | 233 | 4, 518 |
| TCR | C | 1.16e6 | 338 | 5, 1200 | 0.115e6 | 11 | 3, 37 |
| TCR | D | 1.35e6 | 384 | 33, 1401 | 0.116e6 | 11 | 3, 37 |
| TCR | E | 1.98e6 | 1838 | 269, 4828 | 0.350e6 | 127 | 29, 176 |
| TCR | F | 2.98e6 | 2559 | 231, 6561 | 0.813e6 | 294 | 32, 478 |

**FIGURE 2.**
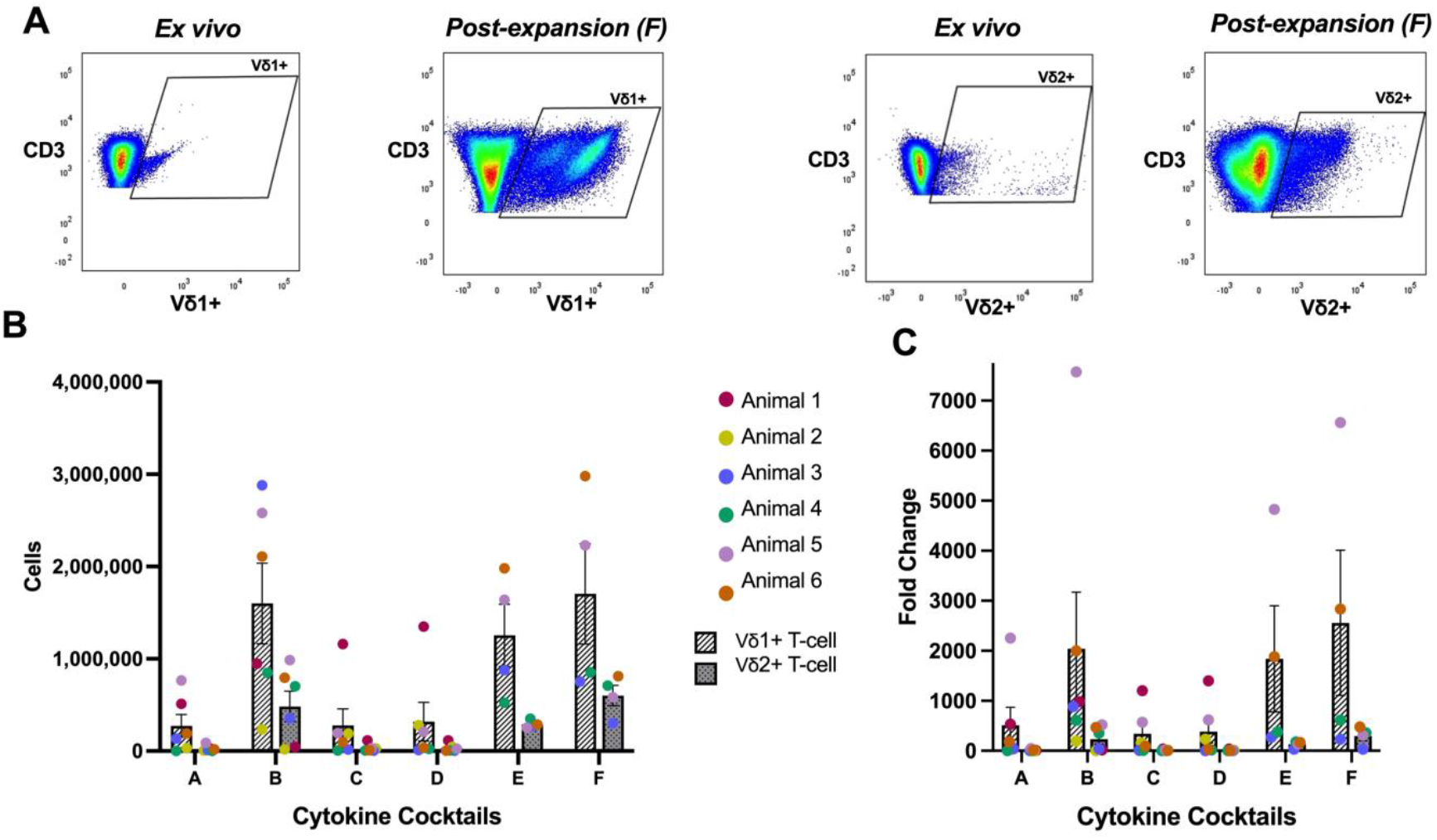
**A**. Representative flow cytometry plots showing ex vivo and post-expansion frequencies of Vδ1^+^ and Vδ2^+^ T cells. **B**. Total Vδ1^+^ and Vδ2^+^ T-cell counts recovered after 9-day expansion of 1×10^6^ PBMCs from Animal 5, demonstrating preferential Vδ1^+^ outgrowth. **C**. Fold change in Vδ1^+^ and Vδ2^+^ T cells after 9 days. The highest average Vδ1^+^ expansion (2,559-fold) occurred with anti-CD3 and cytokine Cocktail F (IL-15 + pVC), with PHA-based Cocktail B yielding similar results (2,040-fold). Data represent mean ± SEM.

Among the tested conditions, the most effective stimulation (CD3 + IL-15 + pVC) achieved up to a 6561-fold expansion (an average of 2559-fold increase). Similarly, the mitogen-based stimulation (PHA + IL-15 + IL-7 + IL-18 + pVC) yielded up to a 7574-fold expansion (an average of 2040-fold increase). Expansions varied among the six animals, with the highest responders starting from the lowest Vδ1^+^ T cell frequencies (0.03% –0.13% of PBMCs) and the lowest responders from higher frequencies (0.19% – 0.5%). While fold changes are influenced by the starting cell frequency, similar trends in absolute cell counts (Fig. 2B) suggest that biological factors, such as competition for space or limiting cytokines, may allow smaller populations to achieve greater proliferative expansion. Across all conditions, Vδ1^+^ T cells consistently exhibited higher expansion than Vδ2^+^ T cells, indicating preferential stimulation of the Vδ1^+^ subset under the tested culture conditions.

### Effect of Cytokine and Stimulatory Cocktails on Vδ1^+^ and Vδ2^+^ T Cell Expansion

We next evaluated the impact of each stimulation and cytokine condition on the fold-expansion within each subset of γδ T cells. Fold-change data for Vδ1^+^ and Vδ2^+^ T cells were log-transformed to facilitate comparison across cytokine stimulation cocktails (Fig. 3). For Vδ1^+^ T cells, significant increases in fold expansion were observed with the addition of pVC (Fig. 3A). Specifically, comparing Cytokine Cocktail A to B (PHA + IL-7, IL-18, IL-15 ± pVC) revealed a significant enhancement of expansion (*p* = 0.0423), demonstrating the beneficial effect of pVC in the PHA-based stimulation. Similarly, the transition from Cytokine Cocktail D to E (α-CD3 + α-CD28 + IL-15 ± pVC) produced a highly significant increase (*p* = 0.0071), confirming a positive effect of pVC in TCR-based stimulation. In contrast, adding IL-2 to the TCR cocktail (C vs. D) did not significantly alter expansion in the absence of pVC. Further, the inclusion of α-CD28 for TCR co-stimulation (E vs. F) showed no significant effect on the fold expansion of Vδ1^+^ T cells.

**FIGURE 3.**
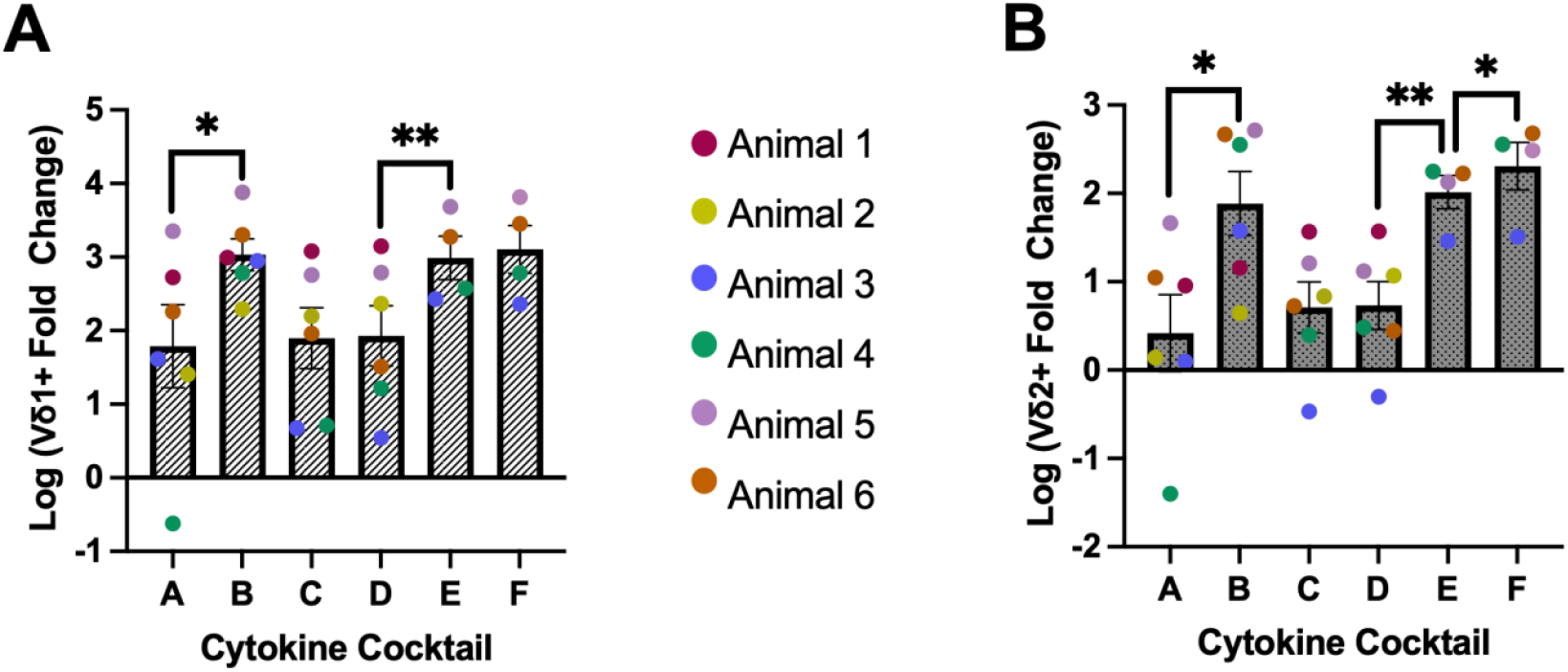
Efficacy of cytokine cocktails for γδ T cell expansion. **A**. Log-transformed Vδ1^+^ fold change after 9-day expansion (skewness: A −1.07, B 0.41, C −0.34, D −0.17, E 0.35, F −0.09). **B**. Log-transformed Vδ2^+^ fold change (skewness: A −0.91, B −0.43, C 0.79, D −0.47, E −1.88, F −1.87). Addition of pVC enhanced fold expansion of both Vδ1^+^ and Vδ2^+^ T cells in PHA- and CD3-based cocktails (B, E, and F). Removal of CD28 (Cocktail E and F) further increased Vδ2^+^ T-cell expansion. Data represent mean ± SEM. Statistical analysis found in Supplemental Materials, Tables 5-6.

For Vδ2^+^ T cells, although the expansion was consistently lower than Vδ1^+^ T cells, the trends were similar, indicating that pVC addition significantly increased Vδ2^+^ expansion in both PHA (A vs. B: *p* = 0.0430) and TCR (D vs. E: *p* = 0.0037) stimulations (Fig. 3B). Impact of a-CD28 in TCR stimulation was found to be significant in Vδ2^+^ T cell expansion, as seen by the significant increase from Cytokine Cocktail E to F (p=0.0438). No significant impact of fold-expansion was observed from Cytokine Cocktail C to D, suggesting no significant impact of IL-2 presence on Vδ2^+^ T cell expansion. These results indicate that pVC consistently enhances overall γδ T cell expansion, whereas IL-2 and α-CD28 have minimal impacts on cell proliferation, suggesting that pVC is the critical component for enhanced γδ T cell expansion in these tested cocktails.

### Quantitative Impact of Cytokine Cocktail Modifications on Vδ1^+^ T Cell Expansion

The efficacy of cytokine cocktail modifications on PHA- and TCR-mediated stimulations was evaluated by calculating the difference in fold change of Vδ1^+^ T cells. Differences were determined by subtracting the average post-expansion fold change of the modified cocktail from that of the standard TCR (α-CD3 + α-CD28 + IL-15) or PHA (PHA + IL-7 + IL-18 + IL-15) expansions. Addition of IL-2 to the standard TCR cocktail (Fig. 4A) resulted in an average decrease in fold change of 46, consistent with the data from Cytokine Cocktails C and D in Fig. 3A, indicating that IL-2 does not significantly enhance Vδ1^+^ T cell expansion in response to TCR stimulation.

**FIGURE 4.**
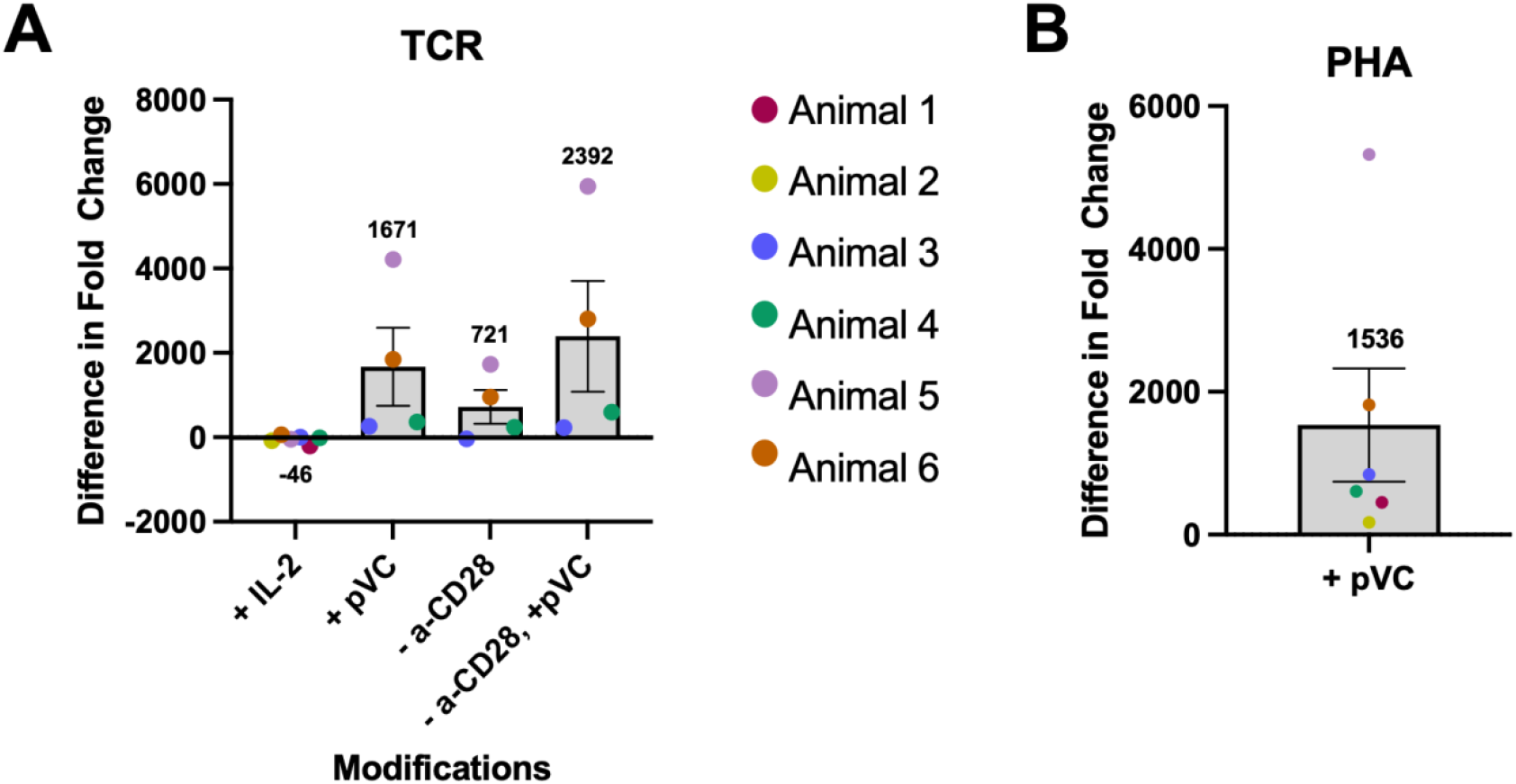
Impact of Cytokine and Costimulatory Modifications on Vδ1^+^ T-Cell Expansion. **A**. Change in Vδ1^+^ fold expansion following stepwise modifications to the CD3/IL-15–based cocktail. Addition of pVC markedly increased Vδ1^+^ expansion (+1671 on average). Removal of anti-CD28 also enhanced expansion (+721), and combining pVC addition with CD28 removal produced the largest gain (+2392). **B**. Change in Vδ1^+^ fold expansion after adding pVC to the PHA/IL-7/IL-18/IL-15 cocktail. Consistent with CD3-based conditions, pVC addition increased Vδ1^+^ expansion (+1536 on average).

In contrast, pVC addition significantly increased Vδ1^+^ T cell expansion for both TCR (Fig. 4A) and PHA (Fig. 4B) stimulations, with average fold-change increases of 1671 and 1536, respectively. Removal of α-CD28 co-stimulation from the TCR cocktail was not statistically significant likely due to donor variability, as seen with the comparison of Cocktails E and F in Fig. 3, but produced an average increase of 721-fold in Vδ1^+^ T cells (Fig. 4A). Notably, combining α-CD28 removal with pVC addition (Cocktail F vs. D) yielded the highest average increase in fold change of 2392, highlighting the compounding effect of these modifications on Vδ1^+^ T cell expansion.

### Expanded Vδ1^+^ T Cells Acquire Effector Memory phenotype

Evaluating the memory phenotype of expanded Vδ1^+^ T cells is crucial for understanding their potential efficacy or in vivo persistence during immunotherapy. Flow cytometric analysis of CD95 versus CD28 expression demonstrated that expanded Vδ1^+^ T cells, when compared to ex vivo donor frequencies, displayed lower frequencies of CD95^+^CD28^+^ central memory cells and higher frequencies of CD95^+^CD28^-^ effector memory subsets (Fig. 5A-C). In vitro expansion resulted in a substantial shift toward a predominant effector memory phenotype, while central memory phenotype was further diminished (Fig. 5B-C) suggesting potential for rapid effector functions and enhanced tissue trafficking. This is consistent with the established effect of IL-15 in driving effector memory phenotype in T cells ^13^. There were no statistically significant changes in the memory phenotype of expanded Vδ1^+^ T cells across all cytokine combinations tested. Overall, these results indicate that expanded Vδ1^+^ T cells exhibit enhanced effector memory features, supporting their functional readiness for use in adoptive immunotherapy applications.

**FIGURE 5.**
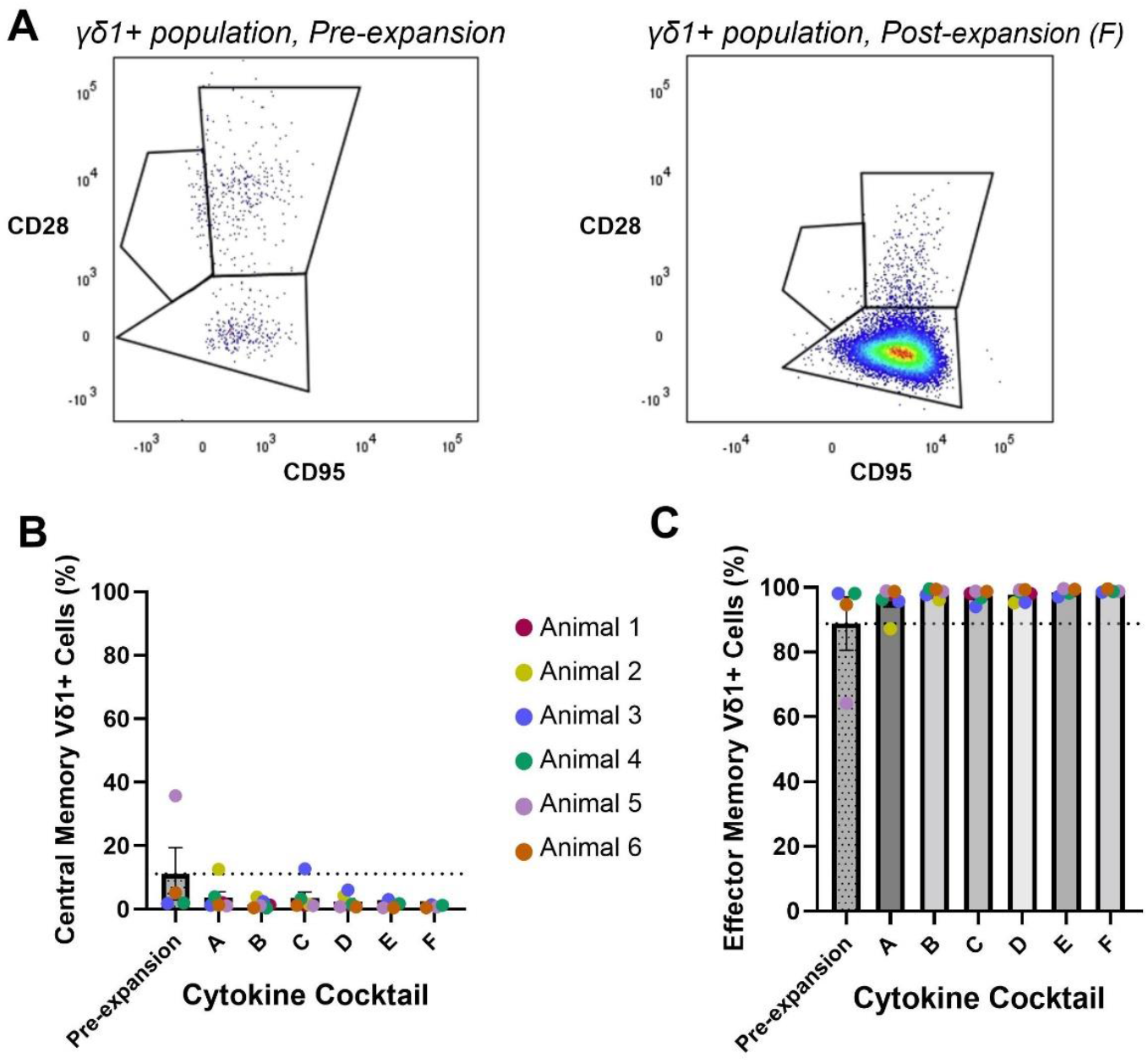
Memory phenotype of expanded Vδ1^+^ T cells. **A**. Representative flow cytometry plots showing central memory (CD28^+^CD95^+^) and effector memory (CD28^-^CD95^+^) subsets within ex vivo and post-expansion Vδ1^+^ T cells from Animal 5. **B**. Frequency of central memory Vδ1^+^ T cells before and after expansion. **C**. Frequency of effector memory Vδ1^+^ T cells before and after expansion. Data represent mean ± SEM. Statistical analysis found in Supplemental Materials, Tables 7-8.

### Distinct Patterns of CD4/CD8 Coreceptor and Transcription Factor Expression in Expanded Vδ1^+^ and Vδ2^+^ T Cells

To further evaluate phenotypic and functional changes during expansion, γδ T cells were immunophenotyped pre- and post-culture based on surface expression of CD3, CD4, CD8, and memory markers along with intracellular staining of lineage-determining transcription factors (LD-TFs) by flow cytometry, allowing delineation of cytokine production and effector polarization ^14^. In vitro expansion amplifies functional heterogeneity within γδ T cells: CD4^+^ cells typically exhibit helper/regulatory attributes, whereas CD8^+^ cells display cytotoxic potential, making coreceptor expression an informative indicator of subset specialization. Consistently, we observed a marked loss of CD4^+^ γδ T cells and maintenance or enrichment of CD8^+^ γδ T cells after 9 days of culture (Fig. 6A–D). For Vδ1^+^ T cells, the predominantly CD8^+^ and CD4^−^/CD8^−^ ex vivo distribution—with minimal CD4 expression—was preserved during expansion (Fig. 6B). In contrast, CD4^+^ Vδ2^+^ T cells declined substantially, from an average of 43.05% pre-expansion to 7.8% post-expansion, accompanied by a shift toward CD8^+^ enrichment, particularly in cultures containing pVC (Fig. 6C–D). Multigraph overlays from a representative macaque demonstrated uniform increases in CD3, CD8, and Vδ1/Vδ2 TCR expression, along with consistent decreases in CD4 and CD28 across media conditions A–F (Fig. 6E–F), confirming acquisition of a CD8-biased effector-memory phenotype.

**FIGURE 6.**
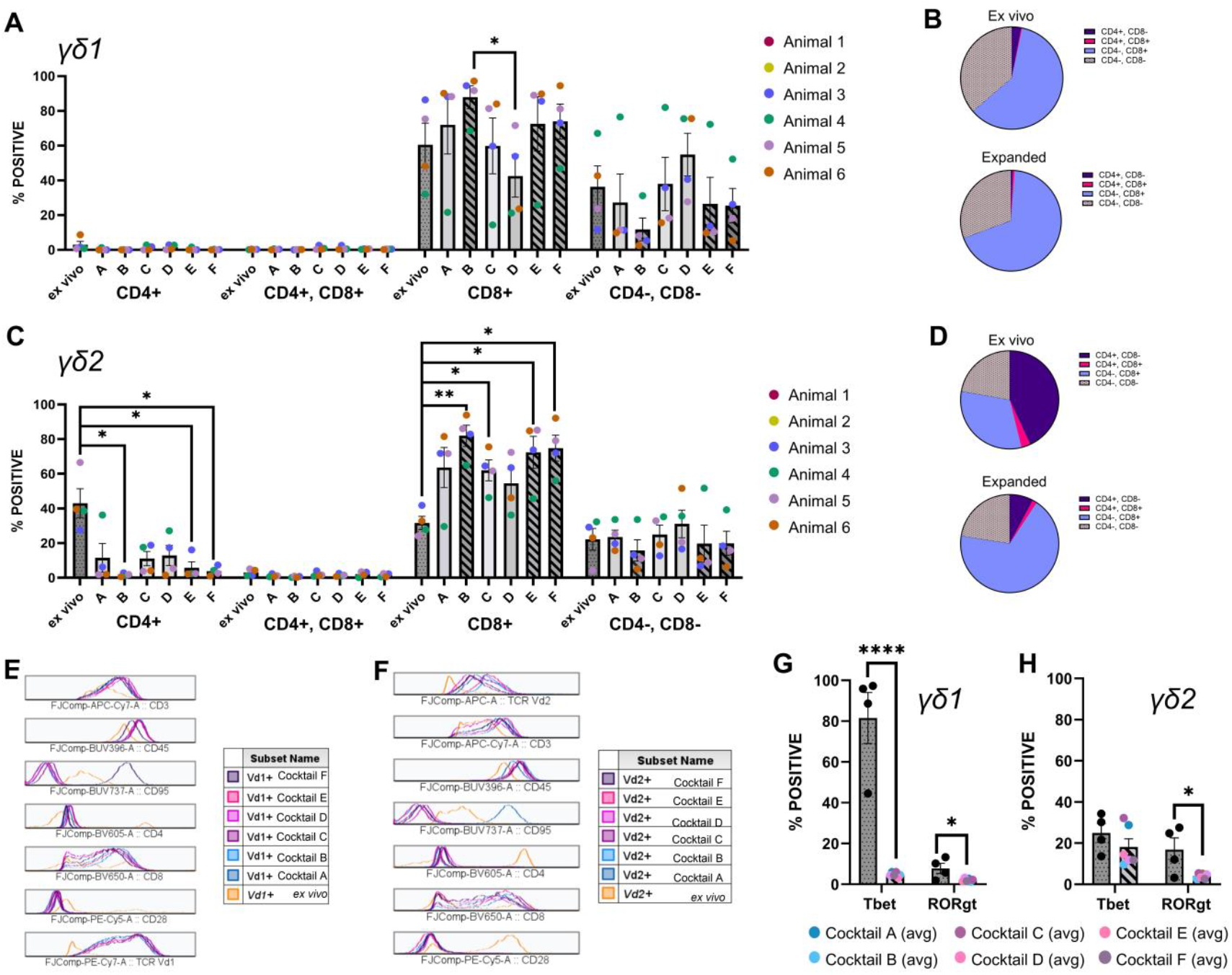
Immunophenotypic characterization of expanded Vδ1^+^ and Vδ2^+^ T cells based on CD4/CD8 Coreceptor and Transcription Factor expression. **A**. Frequencies of CD4^+^, CD4^−^CD8^−^, CD8^+^, and CD4^+^CD8^+^ subsets within Vδ1^+^ T cells ex vivo and following 9-day expansion across all cytokine cocktails (A–F). **B**. Pie charts summarizing the overall distribution of CD4/CD8 coreceptor subsets in ex vivo versus expanded Vδ1^+^ T cells. Statistical Analysis for Figures A-B found in Supplemental Materials, Tables 9-10. **C**. Frequencies of CD4/CD8 subsets within Vδ2^+^ T cells ex vivo and post-expansion. **D**. Pie charts showing the distribution of Vδ2^+^ coreceptor phenotypes before and after culture. **E–F**. Overlay histograms comparing expression of CD3, CD4, CD8, CD28, and Vδ1/Vδ2 TCR in ex vivo versus expanded γδ T cells from a representative animal across cytokine cocktails. **G**. Lineage-determining transcription factor expression in Vδ1^+^ T cells showing a marked reduction in T-bet and RORγt following expansion. **H**. Transcription factor expression in Vδ2^+^ T cells demonstrating maintenance of T-bet with reduced RORγt after expansion. Data represent mean ± SEM; statistical significance determined by paired tests where indicated. Statistical analysis for Figures G-H found in Supplemental Materials, Tables 11-12.

LD-TF analysis revealed that ex vivo Vδ1^+^ T cells expressed markedly higher T-bet (mean 81.6%) and lower RORγt (mean 7.8%), whereas Vδ2^+^ T cells exhibited a more balanced Th1/Th17-like profile with comparable expression of both transcription factors (T-bet mean 24.9%; RORγt mean 16.9%), consistent with our prior findings ^15^. Following expansion, Vδ1^+^ T cells displayed a profound reduction in T-bet (mean 4.8%) alongside a further decrease in RORγt (mean 2.07%) (Fig. 6G). Expanded Vδ2^+^ T cells similarly showed markedly reduced RORγt (mean 4%) while largely maintaining T-bet expression (mean 18.1%) (Fig. 6H). These shifts indicate an overall reduction in T helper-associated transcriptional programming, with more pronounced effects in Vδ1^+^ T cells than in Vδ2^+^ T cells. Collectively, these data indicate that the expansion conditions preferentially drive γδ T cells, particularly Vδ1^+^ cells, toward a CD8-expressing effector-memory phenotype with diminished Th-polarization potential.

## Discussion

Efficient expansion of Vδ1^+^ T cells is a critical prerequisite for advancing their immunotherapeutic application. In this study, we establish a robust macaque PBMC–derived expansion protocol that achieves, to our knowledge, the highest fold increases reported to date. Using CD3 or PHA stimulation in combination with cytokines, including IL-15 and pVC, we generated up to a 7,574-fold increase in Vδ1^+^ T cell numbers within 9 days of culture. The expanded cells acquired a distinct functional profile characterized by diminished Th-polarization and a shift toward a potentially cytotoxic, CD8-expressing effector-memory phenotype in the Vδ1^+^ subset.

Few studies have reported the development of large-scale expansion strategies for Vδ1^+^ T cells. Almeida et al., demonstrated the feasibility of generating cytotoxic, minimally IL-17-producing Vδ1^+^ T cells (DOT cells) through selective cytokine stimulation and anti-CD3 restimulation, achieving up to 2,000-fold expansion of human PBMCs cultured in clinical-grade bags ^9^. Subsequent modifications of this protocol yielded 400-1,000-fold increases in Vδ1^+^ T cells after 20-day culture of αβ- and CD56-depleted and anti-CD3 stimulated PBMCs in the presence of IL-15 ^16^. In comparison, the protocol described here provides substantially higher fold-expansions using a simpler workflow, shorter culture duration, and frozen PBMCs, highlighting its practicality and scalability for Vδ1^+^T cell-specific applications. A more recent evaluation by Verkerk et al. compared PHA stimulation with previously published approaches and demonstrated that both Vδ1^+^ and Vδ2^+^ cytotoxic subsets can be expanded effectively, achieving up to 1,000-fold expansion of Vδ1^+^ T cells ^7^. Although Ferry et al. reported PHA as inferior to anti-CD3 for Vδ1^+^ expansion, Verkerk et al. found that PHA stimulation, specifically with the HA-16 clone, performed robustly when used with feeder cells. Consistent with these findings, our data demonstrated similarly strong Vδ1^+^ T-cell expansion using HA-16 PHA stimulation compared with TCR-driven approaches; however, because IL-7 and IL-18 were included only in the PHA cocktails, differences in expansion may reflect the combined effects of the stimulation mode and these additional cytokines. To ensure consistency across expansion conditions, all cultures used irradiated human PBMCs as feeders; however, this introduces a xenogeneic interaction that likely provides strong accessory co-stimulation and may contribute to the overall magnitude of proliferation observed.

Previous efforts to expand γδ T cells in vitro have relied on diverse combinations of mitogens and cytokines, such as anti-CD3, PHA, zoledronate, concanavalin A, HMBPP, and IL-2, IL-4, IL-7, IL-18, or IL-21, largely optimized for Vδ2^+^ T cells ^17-20^. Far less attention has been given to the role of vitamin C in γδ T-cell culture, despite its known functions in T-cell differentiation, epigenetic regulation, and survival. Based on an earlier study demonstrating enhanced Vδ2^+^ T cell proliferation with pVC ^10^, we explored its effects on Vδ1^+^ T-cell expansion in this study. Across all stimulatory conditions tested, pVC was the single most influential factor determining the magnitude of expansion for both Vδ1^+^ and Vδ2^+^ T cells. Addition of pVC substantially increased fold expansions in both TCR- and mitogen-stimulated PBMC cultures, while IL-2 and CD28 co-stimulation had minimal or no measurable impact. This observation is consistent with broader effects of vitamin C on enhancing in vitro T cell and NK proliferation at optimal concentrations ^21, 22^. Our data extend these findings to Vδ1^+^ γδ T cells, suggesting that pVC may alleviate metabolic bottlenecks associated with rapid proliferation ^23^, enabling up to 7,500-fold increases observed. Moreover, as previously reported, vitamin C can also protect against oxidative stress ^24^, supporting cell survival during periods of rapid immune cell expansion. Previously published literature surrounding Vδ1^+^ T cell expansions in human PBMCs over 10-20 days range from 400 to 2000-fold increases ^7, 10, 11^. The striking effect of pVC in both PHA- and CD3-based conditions in our study supports its incorporation as a critical component for γδ T cell manufacturing protocols.

Unexpectedly, removing CD28 co-stimulation modestly enhanced Vδ1^+^ T cell expansion, and together with pVC, yielded the highest fold increase. This likely reflects lineage-specific requirements since γδ T cells are less dependent on classical CD28 signaling than αβ T cells, which need CD28–CD80/86 engagement for robust proliferation and avoidance of anergy ^25-27^. γδ T cells can proliferate, produce IFN-γ, and exert cytotoxicity without CD28, consistent with their innate-like activation thresholds ^28, 29^. Thus, CD28 stimulation during initial PBMC activation may have preferentially supported αβ T-cell outgrowth, indirectly limiting Vδ1^+^ T cell expansion.

Independent of the stimulation method, Vδ1^+^ T cells consistently expanded more efficiently than Vδ2^+^ T cells, indicating enhanced responsiveness of Vδ1^+^ subsets to IL-15 signaling, which is known to promote homeostatic proliferation and effector memory differentiation, particularly in CD8alpha-expressing cells such as CD8^+^ T cells and NK cells ^13^. These findings align the dominant CD8^+^ phenotype of ex vivo Vδ1^+^ T cells and support previous reports showing that Vδ1^+^ T cells respond robustly to TCR-induced expansion and can be preferentially enriched by cytokine supplementation ^8, 11^. Our results indicate that macaque Vδ1^+^ T cells, similar to their human counterparts, can be selectively expanded ex vivo using defined cocktails, enabling scalable production for downstream immunotherapy applications, including tumor- or viral-targeted adoptive transfer.

Phenotypically, expanded Vδ1^+^ T cells exhibited features of effector-memory differentiation, including loss of CD28, enrichment of CD95^+^CD28^−^ subsets, and maintenance or enhancement of CD8 expression. These characteristics are consistent with IL-15-driven maturation phenotype and may be advantageous for therapeutic use, as effector-memory γδ T cells demonstrate potent cytotoxicity and rapid tissue recruitment ^30, 31^. Importantly, memory differentiation was largely consistent across culture conditions, indicating that the cytokine and stimulatory variations primarily influence proliferative magnitude rather than developmental trajectory. The near-complete loss of CD4 expression and the concurrent enrichment of CD8^+^ γδ T cells further support the emergence of a cytotoxic-biased phenotype. Given that CD4^+^ γδ T cells are typically associated with helper or regulatory functions, their marked reduction during expansion likely reflects a selective advantage for cytotoxic subsets under these culture conditions.

The expanded γδ T cells demonstrated a marked reduction in lineage-defining transcription factors T-bet and RORγt, with the effect especially pronounced in Vδ1^+^ cells. This remodeling suggests a downregulation of polarized Th1/Th17-like transcriptional programs during rapid proliferation. For Vδ2^+^ T cells, T-bet expression was relatively preserved, indicating maintenance of Type 1-associated effector programming, while RORγt decreased substantially. The marked reduction in T-bet in Vδ1^+^ T cells after expansion likely reflects the strong activation and cytokine-driven proliferative environment, rather than functional loss.

This phenomenon is analogous to effector-memory CD8 T^+^ cells, which transiently downregulate T-bet during strong in vivo stimulation to differentiate into memory precursor effector cells capable of generating long-lived memory CD8+ T cells ^32^. Since γδ T cells require acute TCR or stress-ligand engagement to re-express full effector molecules, low T-bet at the end of the culture phase does not necessarily signify impaired cytotoxic potential. These shifts may reflect activation-induced transcriptional resetting, altered cytokine milieu, or selective outgrowth of less polarized subsets during culture. Functionally, reduced LD-TF expression may indicate that expanded γδ T cells adopt a more innate-like effector state, characteristic of many cytotoxic γδ subsets that operate independently of classical Th transcriptional identities. This phenotype aligns well with the enrichment of CD8^+^, effector-memory phenotype cells, and support the concept that expanded Vδ1^+^ T cells remain “poised” for effector function and could re-express functional markers upon restimulation or stress-signal exposure. Future work involving restimulation of the expanded cells and subsequent evaluation of transcription factors like T-bet and Eomesodermin ^33^, alongside key effector molecules, will be essential to prove this concept. This verification is particularly important because a highly cytotoxic and broadly competent phenotype is desirable for γδ T cell-based immunotherapies, where cytotoxic potency and broad cytokine competence are prioritized over helper specialization ^34-37^.

The ability to generate large numbers of macaque Vδ1^+^ T cells with consistent effector-memory phenotypes has important implications for preclinical immunotherapy development. Vδ1^+^ T cells possess potent cytotoxic activity, recognize stress ligands broadly expressed by tumor and infected cells, and exhibit tissue-homing capabilities that may enhance their in vivo effectiveness. Our culture system provides a scalable platform for producing Vδ1^+^ T cells in nonhuman primates, enabling mechanistic evaluation, persistence studies, and efficacy testing in relevant disease models. Culturing of NHP PBMCs with irradiated human PBMC feeder cells provides robust co-stimulation that helps drive the expansion alongside the specific cocktails. Moreover, identifying pVC as a key enhancer of expansion offers a simple, scalable, and cost-effective strategy to improve γδ T cell manufacturing, with clear potential for streamlined clinical-grade production.

Our study has several limitations, including the small sample size, the extremely low ex vivo frequencies of Vδ1^+^ T cells in the study animals, and the substantial inter-individual variability characteristic of γδ T-cell compartments in both macaques and humans. In addition, we did not directly assess the functional output of the expanded Vδ1^+^ T cells, such as cytotoxic granule release, cytokine production, or target-cell killing. Despite these constraints, the robust and reproducible expansion we achieved, together with the clear acquisition of effector-memory phenotypes, establish a framework for deeper functional characterization and further optimization of manufacturing parameters, even for donors with very low circulating Vδ1^+^ T-cell frequencies.

In summary, our findings demonstrate that CD3 or PHA stimulation combined with IL-15 and pVC produces efficient expansion of Vδ1^+^ T cells from macaque PBMCs, yielding phenotypically mature, cytotoxic effector-memory cells with reduced Th polarization. This work defines an optimized protocol for generating large numbers of Vδ1^+^ T cells suitable for mechanistic and translational studies leveraging the unique features of Vδ1^+^ T cells.

## Acknowledgements

We thank the Tulane National Biomedical Research Center (RRID:SCR_008167) and the flow cytometry core (S10 OD026800, RRID: SCR_024611/ SCR_008167) for providing the resources to support this work.

## Data Availability

All data can be found in the manuscript and in the Supplementary Information.

## Author Contributions

IB: Research design, investigation, data analysis and writing (original draft). AM and VH: Research and data analysis. NR: Conceptualization, project administration/supervision and writing (review and editing). YF and NB: Review and editing.

