## Supplemental Data for "Optimized In Vitro Expansion of Vδ1+ γδT Cells from Nonhuman Primate Peripheral Blood"

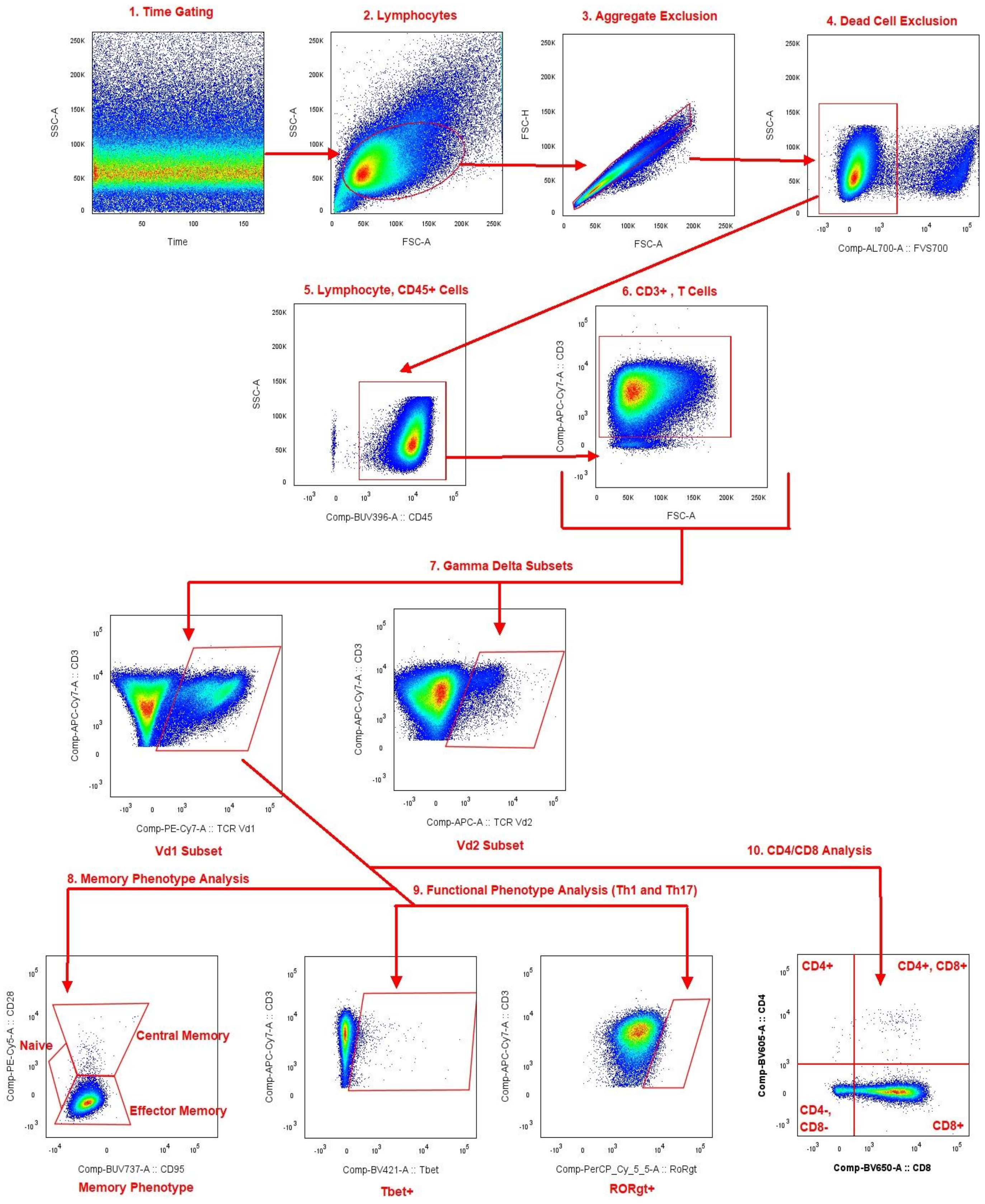


Supplementary Figure 1: Gating Strategy for Flow Cytometry Analysis

Supplementary Table 1: Reagent Supplemental Information

| Reagent | Brand | Catalog No. | Clone |
|---|---|---|---|
| Recombinant Human IL-7 Protein | R&D Systems | Cat# 207-IL-005/CF | |
| Gibco™ Human IL-18BP Fc Recombinant Protein | PeproTech® | Cat# 20018BP20UG | |
| Human IL-2 IS premium grade | Miltenyi Biotec | Cat# 130-097-748 | |
| Recombinant Human IL-15 carrier-free | Biolegend | Cat# 570306 | |
| phospho-Vitamin C | Millipore Sigma | Cat# A8960-5G | |
| PHA Purified | ThermoScientific | Cat# R30852801 | HA16 |
| a-CD3 | NHP Reagent Resource | RRID:AB_2819276 | FN18 |
| a-CD28 | NHP Reagent Resource | RRID: AB_2819274 | a-CD28.2 |

Supplementary Table 2: Cytokine Cocktails– 1 mL total per well, no mitogen restimulation, 1e6 seeding rate of NHP PBMCs with 1:2 ratio of NHP PBMCs: human irradiated feeder cells.

| Cocktail | Cytokine concentrations per well | | | | |
|---|---|---|---|---|---|
| A | PHA | IL-7 | IL-18 | IL-15 | |
| | 1 ug/mL | 20 ng/mL | 10 ng/mL | 20 ng/mL | |
| B | PHA | IL-7 | IL-18 | IL-15 | Vit C |
| | 1 ug/mL | 20 ng/mL | 10 ng/mL | 20 ng/mL | 50 ug/mL |
| C | CD3 | CD28 | IL-2 | IL-15 | |
| | 5 ug/mL | 2.5 ug/mL | 50 IU/mL | 20 ng/mL | |
| D | CD3 | CD28 | IL-15 | | |
| | 5 ug/mL | 2.5 ug/mL | 20 ng/mL | | |
| E | CD3 | CD28 | IL-15 | Vit C | |
| | 5 ug/mL | 2.5 ug/mL | 20 ng/mL | 50 ug/mL | |
| F | CD3 | IL-15 | Vit C | | |
| | 5 ug/mL | 20 ng/mL | 50 ug/mL | | |

Supplementary Table 3: Antibody Supplemental Information for Flow Cytometry

| Antibody | Channel | Brand | Catalog No. | Clone |
|---|---|---|---|---|
| TCR γ9 | FITC | Invitrogen | Cat# TCR2720 | 7A5 |
| RORγt | PCP-Cy5.5 | Invitrogen | Cat# 46-6988-82 | AFKJS-9 |
| T-bet | BV421 | Biolegend | Cat# 644816 | 4B10 |
| TCR γδ | BV510 | Biolegend | Cat# 331220 | B1 |
| CD4 | BV605 | BD Horizon | Cat# 562843 | L200 |
| CD8 | BV650 | Bd Horizon, | Cat #565289 | SK1 |
| CD45 | BUV395 | BD Horizon | Cat# 564099 | D058-1283 |
| CD95 | BUV737 | BD Horizons | Cat# 612790 | DX2 |
| TCR γ4 | PE | Invitrogen | Cat# MA5-44121 | 4A11.904 |
| AhR | PE-CF549 | BD Horizon | Cat# 565790 | T49-550 |

Supplementary Table 4: Supplemental Expansion Data of Vδ1+ T Cells after Expansion

| **Stimulation** | **Cocktail** | **Average Viability of PBMCs(%) after 9-day culture** | **Avg Purity of Vδ1+ T cells (from CD3+ parent)** | **Avg Purity of Vδ2+ T cells (from CD3+ parent)** |
|---|---|---|---|---|
| PHA | A | 31.80% | 32.7% | 3.45% |
| PHA | B | 60.63% | 35.2% | 9.41% |
| TCR | C | 23.39% | 25.1% | 4.74% |
| TCR | D | 23.74% | 22.8% | 4.32% |
| TCR | E | 70.93% | 14.8% | 3.58% |
| TCR | F | 72.23% | 16.9% | 6.67% |

Supplementary Table 5: Statistical Analysis of Expansion *Figure 3a*, Log (Vδ1+ Fold Change) after 9-day Expansion

| Comparison | Statistical Comparison | P-value | Significance | t-score | n |
|---|---|---|---|---|---|
| +/- pVC in PHA stimulation | Cytokine Cocktail A vs. B | 0.0423 | * | 2.710 | 6 |
| +/- IL-2 in TCR stimulation | Cytokine Cocktail C vs. D | 0.8073 | ns | 0.2572 | 6 |
| +/- pVC in TCR stimulation | Cytokine Cocktail D vs. E | 0.0071 | ** | 6.585 | 4 |
| +/- CD28 in TCR stimulation with pVC | Cytokine Cocktail E vs. F | 0.1630 | ns | 1.840 | 4 |

Supplementary Table 6: Statistical Analysis of Expansion *Figure 3b*, Log (Vδ2+ Fold Change) after 9-day Expansion

| Comparison | Statistical Comparison | P-value | Significance | t-score | n |
|---|---|---|---|---|---|
| +/- pVC in PHA stimulation | Cytokine Cocktail A vs. B | 0.0430 | * | 2.695 | 6 |
| +/- IL-2 in TCR stimulation | Cytokine Cocktail C vs. D | 0.7931 | ns | 0.2767 | 6 |
| +/- pVC in TCR stimulation | Cytokine Cocktail D vs. E | 0.0037 | ** | 8.263 | 4 |
| +/- CD28 in TCR stimulation with pVC | Cytokine Cocktail E vs. F | 0.0438 | * | 3.358 | 4 |

Supplementary Table 7: Statistical Analysis of Expansion *Figure 4b*, Percent of Central Memory Phenotypes Vδ1⁺ T-cells in pre-expanded PBMCs and in Vδ1⁺ T-cell expansion cultures.

| Statistical Comparison | P-value | Significance | t-score | n |
|---|---|---|---|---|
| Cytokine Cocktail A vs. Pre-expansion | 0.3561 | ns | 1.088 | 4 |
| Cytokine Cocktail B vs. Pre-expansion | 0.3100 | ns | 1.219 | 4 |
| Cytokine Cocktail C vs. Pre-expansion | 0.5527 | ns | 0.666 | 4 |
| Cytokine Cocktail D vs. Pre-expansion | 0.3895 | ns | 1.004 | 4 |
| Cytokine Cocktail E vs. Pre-expansion | 0.3406 | ns | 1.130 | 4 |
| Cytokine Cocktail F vs. Pre-expansion | 0.2997 | ns | 1.251 | 4 |

Supplementary Table 8: Statistical Analysis of Expansion *Figure 4c*, Percent of Effector Memory Phenotypes Vδ1⁺ T-cells in pre-expanded PBMCs and in Vδ1⁺ T-cell expansion cultures.

| Statistical Comparison | P-value | Significance | t-score | n |
|---|---|---|---|---|
| Cytokine Cocktail A vs. Pre-expansion | 0.3921 | ns | 0.9973 | 4 |
| Cytokine Cocktail B vs. Pre-expansion | 0.3174 | ns | 1.197 | 4 |
| Cytokine Cocktail C vs. Pre-expansion | 0.4125 | ns | 0.950 | 4 |
| Cytokine Cocktail D vs. Pre-expansion | 0.3557 | ns | 1.089 | 4 |
| Cytokine Cocktail E vs. Pre-expansion | 0.3315 | ns | 1.156 | 4 |
| Cytokine Cocktail F vs. Pre-expansion | 0.3170 | ns | 1.198 | 4 |

Supplementary Table 9: Statistical Analysis of Expansion *Figure 6a*, Frequencies of $CD4^{+}$, $CD4^{-}CD8^{-}$, $CD8^{+}$, and $CD4^{+}CD8^{+}$ subsets within $V\delta1^{+}$ T cells ex vivo and following 9-day expansion across all cytokine cocktails (A–F).

| Statistical Comparison | P-value | Significance | t-score | n |
|---|---|---|---|---|
| CD4+: Cocktail A vs. Ex vivo | 0.2959 | ns | 1.263 | 4 |
| CD4+: Cocktail B vs. Ex vivo | 0.2270 | ns | 1.515 | 4 |
| CD4+: Cocktail C vs. Ex vivo | 0.5045 | ns | 0.7562 | 4 |
| CD4+: Cocktail D vs. Ex vivo | 0.5926 | ns | 0.5970 | 4 |
| CD4+: Cocktail E vs. Ex vivo | 0.3188 | ns | 1.192 | 4 |
| CD4+: Cocktail F vs. Ex vivo | 0.2533 | ns | 1.410 | 4 |
| CD4+, CD8+: Cocktail A vs. Ex vivo | 0.8428 | ns | 0.2161 | 4 |
| CD4+, CD8+: Cocktail B vs. Ex vivo | 0.1469 | ns | 1.945 | 4 |
| CD4+, CD8+: Cocktail C vs. Ex vivo | 0.2774 | ns | 1.324 | 4 |
| CD4+, CD8+: Cocktail D vs. Ex vivo | 0.3003 | ns | 1.249 | 4 |
| CD4+, CD8+: Cocktail E vs. Ex vivo | 0.1525 | ns | 1.907 | 4 |
| CD4+, CD8+: Cocktail F vs. Ex vivo | 0.4550 | ns | 0.8559 | 4 |
| CD8+: Cocktail A vs. Ex vivo | 0.3775 | ns | 1.033 | 4 |
| CD8+: Cocktail B vs. Ex vivo | 0.0603 | ns | 2.944 | 4 |
| CD8+: Cocktail C vs. Ex vivo | 0.9671 | ns | 0.0448 | 4 |
| CD8+: Cocktail D vs. Ex vivo | 0.0703 | ns | 2.757 | 4 |
| CD8+: Cocktail E vs. Ex vivo | 0.3420 | ns | 1.126 | 4 |
| CD8+: Cocktail F vs. Ex vivo | 0.3593 | ns | 1.080 | 4 |
| CD4-, CD8-: Cocktail A vs. Ex vivo | 0.3975 | ns | 0.9845 | 4 |
| CD4-, CD8-: Cocktail B vs. Ex vivo | 0.0567 | ns | 3.021 | 4 |
| CD4-, CD8-: Cocktail C vs. Ex vivo | 0.8941 | ns | 0.1447 | 4 |
| CD4-, CD8-: Cocktail D vs. Ex vivo | 0.0829 | ns | 2.564 | 4 |
| CD4-, CD8-: Cocktail B vs. Cocktail D | 0.0313 | * | 3.835 | 4 |
| CD4-, CD8-: Cocktail E vs. Ex vivo | 0.3445 | ns | 1.120 | 4 |
| CD4-, CD8-: Cocktail F vs. Ex vivo | 0.3915 | ns | 0.9988 | 4 |

Supplementary Table 10: Statistical Analysis of Expansion *Figure 6c*, Frequencies of CD4/CD8 subsets within Vδ2$^{+}$ T cells ex vivo and post-expansion.

| Statistical Comparison | P-value | Significance | t-score | n |
|---|---|---|---|---|
| CD4+: Cocktail A vs. Ex vivo | 0.0969 | ns | 2.388 | 4 |
| CD4+: Cocktail B vs. Ex vivo | 0.0159 | * | 4.944 | 4 |
| CD4+: Cocktail C vs. Ex vivo | 0.0706 | ns | 2.753 | 4 |
| CD4+: Cocktail D vs. Ex vivo | 0.0889 | ns | 2.485 | 4 |
| CD4+: Cocktail E vs. Ex vivo | 0.0405 | * | 3.464 | 4 |
| CD4+: Cocktail F vs. Ex vivo | 0.0231 | * | 4.303 | 4 |
| CD4+, CD8+: Cocktail A vs. Ex vivo | 0.2232 | ns | 1.531 | 4 |
| CD4+, CD8+: Cocktail B vs. Ex vivo | 0.0822 | ns | 2.575 | 4 |
| CD4+, CD8+: Cocktail C vs. Ex vivo | 0.5010 | ns | 0.7629 | 4 |
| CD4+, CD8+: Cocktail D vs. Ex vivo | 0.2803 | ns | 1.314 | 4 |
| CD4+, CD8+: Cocktail E vs. Ex vivo | 0.4029 | ns | 0.9716 | 4 |
| CD4+, CD8+: Cocktail F vs. Ex vivo | 0.2263 | ns | 1.518 | 4 |
| CD8+: Cocktail A vs. Ex vivo | 0.0612 | ns | 2.925 | 4 |
| CD8+: Cocktail B vs. Ex vivo | 0.0045 | ** | 7.715 | 4 |
| CD8+: Cocktail C vs. Ex vivo | 0.0136 | * | 5.235 | 4 |
| CD8+: Cocktail D vs. Ex vivo | 0.0819 | ns | 2.578 | 4 |
| CD8+: Cocktail E vs. Ex vivo | 0.0249 | * | 4.182 | 4 |
| CD8+: Cocktail F vs. Ex vivo | 0.0130 | * | 5.313 | 4 |
| CD4-, CD8-: Cocktail A vs. Ex vivo | 0.8600 | ns | 0.1921 | 4 |
| CD4-, CD8-: Cocktail B vs. Ex vivo | 0.3752 | ns | 1.039 | 4 |
| CD4-, CD8-: Cocktail C vs. Ex vivo | 0.7908 | ns | 0.2899 | 4 |
| CD4-, CD8-: Cocktail D vs. Ex vivo | 0.3880 | ns | 1.007 | 4 |
| CD4-, CD8-: Cocktail E vs. Ex vivo | 0.8005 | ns | 0.2760 | 4 |
| CD4-, CD8-: Cocktail F vs. Ex vivo | 0.7569 | ns | 0.3391 | 4 |

Supplementary Table 11: Statistical Analysis of *Figure 6g*, Lineage-determining transcription factor expression in Vδ1$^{+}$ T cells showing a marked reduction in T-bet and RORγt following expansion.

| Statistical Comparison | P-value | Significance | t-score | n |
|---|---|---|---|---|
| Tbet+: Ex vivo vs. Expanded | <0.0001 | **** | 7.756 | 4 |
| RORgt+: Ex vivo vs. Expanded | 0.0224 | * | 2.823 | 4 |

Supplementary Table 12: Statistical Analysis of *Figure 6h*, Transcription factor expression in Vδ2$^{+}$ T cells demonstrating maintenance of T-bet with reduced RORγt after expansion.

| Statistical Comparison | P-value | Significance | t-score | n |
|---|---|---|---|---|
| Tbet+: Ex vivo vs. Expanded | 0.2998 | ns | 1.109 | 4 |
| RORgt+: Ex vivo vs. Expanded | 0.0215 | * | 2.849 | 4 |
